# Preoperative predictions of in-hospital mortality using electronic medical record data

**DOI:** 10.1101/329813

**Authors:** Brian Hill, Robert Brown, Eilon Gabel, Christine Lee, Maxime Cannesson, Loes Olde Loohuis, Ruth Johnson, Brandon Jew, Uri Maoz, Aman Mahajan, Sriram Sankararaman, Ira Hofer, Eran Halperin

## Abstract

**Background:** Predicting preoperative in-hospital mortality using readily-available electronic medical record (EMR) data can aid clinicians in accurately and rapidly determining surgical risk. While previous work has shown that the American Society of Anesthesiologists (ASA) Physical Status Classification is a useful, though subjective, feature for predicting surgical outcomes, obtaining this classification requires a clinician to review the patient’s medical records. Our goal here is to create an improved risk score using electronic medical records and demonstrate its utility in predicting in-hospital mortality without requiring clinician-derived ASA scores.

**Methods:** Data from 49,513 surgical patients were used to train logistic regression, random forest, and gradient boosted tree classifiers for predicting in-hospital mortality. The features used are readily available before surgery from EMR databases. A gradient boosted tree regression model was trained to impute the ASA Physical Status Classification, and this new, imputed score was included as an additional feature to preoperatively predict in-hospital post-surgical mortality. The preoperative risk prediction was then used as an input feature to a deep neural network (DNN), along with intraoperative features, to predict postoperative in-hospital mortality risk. Performance was measured using the area under the receiver operating characteristic (ROC) curve (AUC).

**Results:** We found that the random forest classifier (AUC 0.921, 95%CI 0.908-0.934) outperforms logistic regression (AUC 0.871, 95%CI 0.841-0.900) and gradient boosted trees (AUC 0.897, 95%CI 0.881-0.912) in predicting in-hospital post-surgical mortality. Using logistic regression, the ASA Physical Status Classification score alone had an AUC of 0.865 (95%CI 0.848-0.882). Adding preoperative features to the ASA Physical Status Classification improved the random forest AUC to 0.929 (95%CI 0.915-0.943). Using only automatically obtained preoperative features with no clinician intervention, we found that the random forest model achieved an AUC of 0.921 (95%CI 0.908-0.934). Integrating the preoperative risk prediction into the DNN for postoperative risk prediction results in an AUC of 0.924 (95%CI 0.905-0.941), and with both a preoperative and postoperative risk score for each patient, we were able to show that the mortality risk changes over time.

**Conclusions:** Features easily extracted from EMR data can be used to preoperatively predict the risk of in-hospital post-surgical mortality in a fully automated fashion, with accuracy comparable to models trained on features that require clinical expertise. This preoperative risk score can then be compared to the postoperative risk score to show that the risk changes, and therefore should be monitored longitudinally over time.

**Author summary:** Rapid, preoperative identification of those patients at highest risk for medical complications is necessary to ensure that limited infrastructure and human resources are directed towards those most likely to benefit. Existing risk scores either lack specificity at the patient level, or utilize the American Society of Anesthesiologists (ASA) physical status classification, which requires a clinician to review the chart. In this manuscript we report on using machine-learning algorithms, specifically random forest, to create a fully automated score that predicts preoperative in-hospital mortality based solely on structured data available at the time of surgery. This score has a higher AUC than both the ASA physical status score and the Charlson comorbidity score. Additionally, we integrate this score with a previously published postoperative score to demonstrate the extent to which patient risk changes during the perioperative period.

## Introduction

A small number of high-risk patients comprise the majority of patients with surgical complications [1]. Furthermore, many studies have demonstrated that early interventions can help reduce or even prevent perioperative complications [2, 3]. Thus, in the current value-based care environment, it is critical to have methods to rapidly identify those patients who are at the highest risk and thus most likely to benefit from labor or cost-intensive interventions. Unfortunately, many current methods of risk stratification either lack precision on a patient level or require a trained clinician to review the medical records.

Existing preoperative patient risk scores fall into one of two groups. Some attempt to leverage International Statistical Classification of Diseases and Related Health Problems (ICD) codes in order to create models of risk [4–6]. Unfortunately, ICD codes are not available until after patient discharge and have been repeatedly shown to lack accuracy at the patient level [7]. Thus, while these scores tend to perform well for populations, they do not rely on data available immediately prior to surgery and they ultimately lack precision. The other group of models rely on subjective clinician judgment, seen with the American Society of Anesthesiologists Physical Status Score (ASA Score) alone or when incorporated into another model [8]. While these scores tend to have increased precision compared to ICD codes, they cannot be fully automated due to the need for a highly trained clinician to manually review the patient’s chart prior to calculation.

Recently, attempts have been made to leverage machine learning techniques using healthcare data in order to improve the predictive ability of various models [9, 10]. These methods have shown progress in leveraging increasingly complex data while still allowing for the full automation of the scoring system.

In this manuscript, we describe a random forest-based model for the prediction of in-hospital post-surgical mortality using only EMR features readily available before surgery that require no intervention by clinical staff. We compare the performance of our model to existing clinical risk scores (ASA Score and Charlson Comorbidity Score [6]) and show that our model has a considerably higher area under the curve when compared to these. Last, we integrate our model with a previously published model [11] to automatically predict in-hospital mortality immediately after surgery in order to quantify the change in risk during the perioperative period, and we show that, while the majority of patients do not exhibit a substantial change in risk, a subset of the population does show a large change in risk and, therefore, the risk should be monitored at multiple time points.

## 1 Materials and methods

### 1.1 Data source and extraction

All data for this study were extracted from the Perioperative Data Warehouse (PDW), a custom built, robust data warehouse containing all patients who have undergone surgery at UCLA Health since the implementation of our EMR (EPIC Systems, Madison, WI) in March 2013. We have previously described the creation of the PDW, which has a two stage design [12]. Briefly, in the first stage, data are extracted from EPIC’s Clarity database into 26 tables organized around three distinct concepts: patients, surgical procedures and health system encounters. These data are then used to populate a series of 800 distinct measures and metrics such as procedure duration, readmissions, admission ICD codes, and others. All data used for this study were obtained from this data warehouse and an IRB approval (IRB#16-001768) was obtained with exemption status for this retrospective review.

### 1.2 Model Endpoint Definition

We trained classification models to predict in-hospital mortality as a binary outcome. This classification was extracted from the PDW and was set to one if a “death date” was noted during the hospitalization, or the final disposition was set to expired and there were no future admissions for the patient and a clinician “death note” existed. Due to the concern about the need to eliminate false positive results, the results of this definition were validated by trained clinicians in a subset of patients.

### 1.3 Electronic Medical Record (EMR) Data Filtering

A number of filtering criteria were applied before conducting our analysis, as shown by the CONSORT diagram in Fig 1. If a patient underwent multiple surgeries in a single admission, only the first surgical operation was used. Only surgical patients undergoing general anesthesia were considered, as documented by the anesthesiologist note, since we were interested in surgeries with relatively high risk. Any patients that did not have data for the outcome of interest in the database (i.e. had not yet been discharged) were removed from the cohort. Outpatient surgeries were filtered out, and cases with an ASA Physical Status of 6 (organ donors) were also excluded. Patients younger than 18 years old were excluded and anyone older than 89 years old were removed due to HIPAA-based privacy concerns.

**Fig 1.**
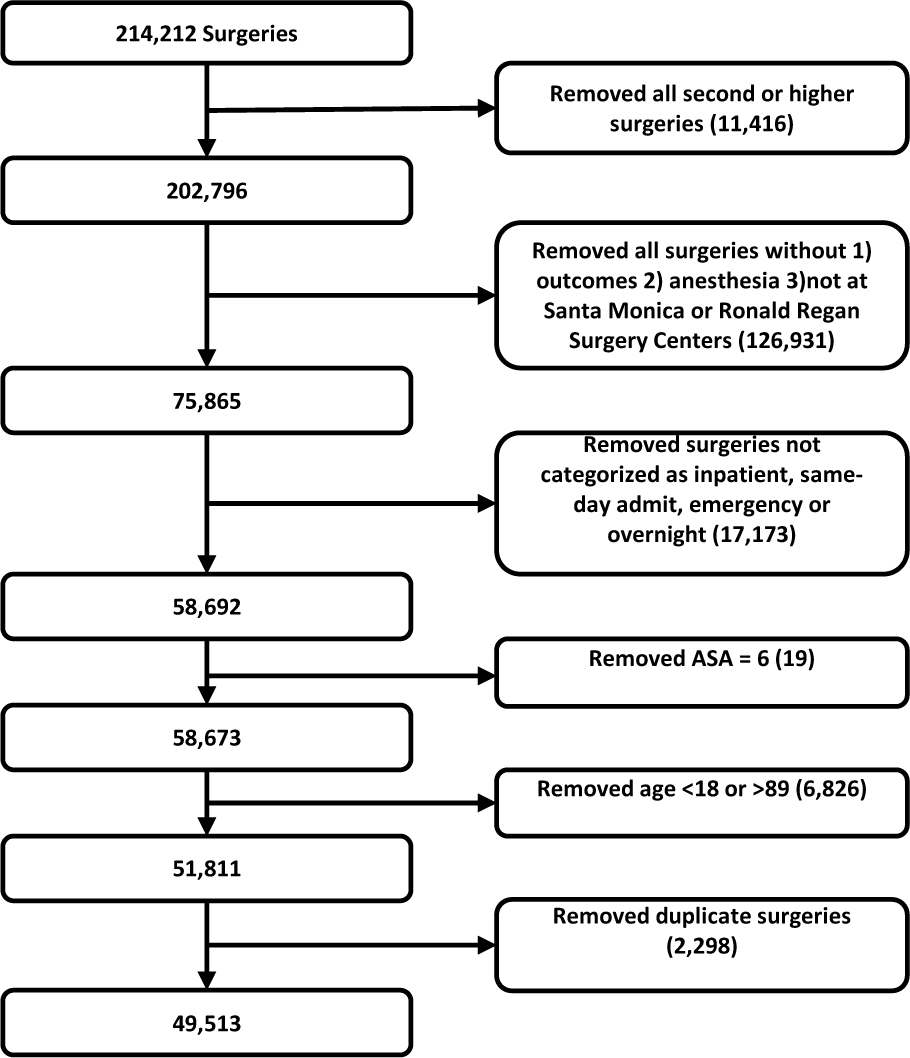
Consort Diagram. Filtering steps taken to select our cohort.

### 1.4 Model Input Features

We tested model performance using seven different sets of input features: (1) Charlson comorbidity score [6] as the only input feature, (2) ASA Physical Status as the only input feature, (3) preoperative features without the ASA status, (4) preoperative features and the ASA status, (5) preoperative features with the imputed-ASA score (see Section 1.6), (6) preoperative features and the ASA status, excluding variables indicating how long before surgery each lab is time-stamped as having been run (not ordered), and (7) preoperative features and the imputed-ASA, excluding variables indicating how long before surgery each lab is time-stamped. The lab result time-stamps were removed from feature sets 6 and 7 because, while they may be correlated with negative outcomes, they are not a direct reflection of patient health. All input features, with the exception of the ASA Physical Status, are accessible automatically before surgery via the EMR system (and were extracted here by the PDW). The ASA Physical Status is manually entered into the system by the anesthesiologist after reviewing the patient’s record. Model features include basic patient information such as age, sex, BMI, blood pressure, and pulse, lab tests frequently given prior to surgery such as sodium, potassium, creatinine, and blood cell counts, and surgery specific information such as the surgical codes. In total, 35 preoperative features (including ASA status) were used and a full list is available in Supplemental Table S1.

### 1.5 Data Preprocessing

Data points greater than 4 standard deviations from the mean were removed. Missing data were imputed using the SoftImpute algorithm [13], as implemented by the fancyimpute Python package [14], with a maximum of 200 iterations. Categorical features were converted into indicator variables, and the first variable was dropped. As the number of positive training examples (in-patient mortalities) was much smaller than the number of negative training examples (survivors), the class balance was greatly skewed (0.61% mortality rate). To overcome this issue, each training set in the cross-validation was oversampled using the SMOTE algorithm [15], implemented in the imblearn Python package [16], using 3 nearest neighbors and the “baseline1” method to create a balanced class distribution. The testing sets were not oversampled and, therefore, maintained the natural outcome frequency. Finally, the training data features were rescaled to have a mean of 0 and standard deviation of 1, and the test data were rescaled using the training data means and standard deviations.

### 1.6 Generating imputed-ASA scores using ASA Status

The ASA Physical Status Classification score is a subjective assessment of a patient’s overall health [17, 18]. The score can take on one of six values, with a score of 1 being a healthy patient and 6 being a brain-dead organ donor [19]. An ASA Physical Status Classification is commonly used in two ways: as a quantitative measure of patient health before a surgery, and for healthcare billing [19]. The ASA Physical Status Classification has been shown to be a useful feature in predicting surgical outcomes [17, 20–22]. However, its inter-rater reliability, a measure of concurrence between scores given by different individuals, is moderate [18].

While the ASA status is a strong predictor of patient health status [17, 20–22], this classification requires a clinician to look through the patient’s chart and subjectively determine the score, which consumes valuable time and requires clinical expertise. In order to balance the value of this score with the desire for automation, we sought to generate a similar metric using readily available data from the EMR - an “imputed” ASA score. Recent works have similarly attempted to develop machine learning approaches to predict ASA scores [23, 24]. However, these methods have difficulty classifying ASA scores of 4 and 5 due to the low frequency of occurrence, and resort to either grouping classes together or ignoring patients with an ASA status of 5. The goal in our work is not to supplant the ASA score but rather to estimate a measure of general patient health for use in our model without needing the time-consuming clinician chart review.

Using the existing ASA Physical Status Classification extracted from the EMR data, we trained a gradient boosted tree regression model to predict the ASA status of new patients using preoperative features unrelated to the surgery. The model was implemented using the XGBoost package [25] with 2000 trees and a max tree depth of 7. We used 10-fold cross-validation to generate predictions. This imputed-ASA value is a continuous number, unlike the actual ASA status which is limited to integers. We compared the performance of a model that uses the imputed-ASA scores and preoperative features to a model that uses the actual ASA Physical Status values and preoperative features.

### 1.7 Model Testing and Training

We evaluated four different classification models: logistic regression with two different types of regularization, random forest classifiers, and gradient boosted tree classifiers. Logistic regression classifiers were trained with both an L2 penalty and an ElasticNet [26] penalty, where alpha (regularization constant) and the L1/L2 mixing parameter were set using 5-fold cross-validation. The random forest classifiers were trained with 2000 estimators, Gini impurity as the splitting criterion, and no maximum tree depth was specified. The gradient boosted machine classifiers were trained using 2000 estimators and a max tree depth of 5. All predictions were made using 10-fold cross-validation on the entire dataset. The logistic regression and random forest classifiers were implemented using Scikit-learn [27], and the gradient boosted tree classifiers were implemented using the XGBoost package [25]. All performance metrics were calculated using methods implemented by Scikit-learn [27].

### 1.8 Feature Importance

To determine which features were most important to the classification models, we examined the model weights for linear models, the feature (Gini) importance for the random forest models, and the feature weight (number of times a feature appears in a tree) for the gradient boosted tree models.

### 1.9 Integrating Preoperative Risk with Postoperative Risk

Previous work [11] has shown that integrating a measure of preoperative risk into a postoperative mortality risk prediction model increases the model performance. We integrated the risk predictions of our random forest model with the previously developed postoperative model in order to see if our model predictions can similarly serve this function. Using the deep neural network architecture and features used by Lee et al. [11], we replaced the ASA status feature with the preoperative risk score (generated using the random forest model with preoperative features and imputed-ASA scores) and trained the model on the same cohort used for preoperative risk prediction. The intraoperative data were preprocessed in the same manner as described in [11]. We then compared the area under the ROC of the postoperative model trained using the ASA status and intraoperative features to the postoperative model that was trained using the preoperative risk score and intraoperative features.

### 1.10 Generating Charlson Comorbidity Indexes

We compared our method with the Charlson Comorbidity Index scores [6], a well-known and proven existing method for prediction of risk of postoperative mortality, for each patient in the cohort. We used the updated weights as described by Quan et al. [28]. Scores were calculated using the R package *icd* [29] on all ICD10 codes associated with each surgery admission.

## 2 Results

### 2.1 Patient Demographics

The patient dataset contained 49,513 surgical records (see Table 1). These patients were between the ages of 18 and 89, with a mean age of 56, and were treated as either inpatients, same-day admits, emergencies, or overnight recoveries. The frequency of mortality in the dataset was approximately 0.61%. A patient ASA status of 3 was the most common, comprising 47.5% of the dataset.

**Table 1.** Patient Demographics.

| Property | Quantification |
| --- | --- |
| Number of Patients | 49,513 |
| Patients with in-hospital mortality | 301 (0.61%) |
| Mean Age | 56.0 (s.d. 17.3) |
| Female patients | 25,983 (52.5%) |
| ASA status - 1 | 3,479 (7.0%) |
| ASA status - 2 | 19,589 (39.6%) |
| ASA status - 3 | 23,543 (47.5%) |
| ASA status - 4 | 2,775 (5.6%) |
| ASA status - 5 | 127 (0.3%) |
| Ronald Reagan Surgical Center | 31835 (64.3%) |
| Santa Monica Surgical Center | 17678 (35.7%) |

Patient demographics for the cohort used for training and testing models. Number of patients and percent of the total cohort are shown.

### 2.2 Model Performance

#### 2.2.1 Area under the receiver operating characteristic (ROC) curve

The area under the ROC curve values for each model are shown in Table 2 and ROC curves are shown for the random forest model in Fig 2a and for all models in Supplemental Fig S1. Models using the preoperative features have higher area under the ROC values (0.909, 95%CI 0.881-0.937) than the models that use either the Charlson comorbidity score (0.706, 95%CI 0.669-0.744) or the ASA status (0.865, 95%CI 0.848-0.882). Adding the imputed-ASA status values to the preoperative features improves the area under the ROC (0.912, 95%CI 0.902-0.923), but using the true ASA value assigned by anesthesiologists, along with the preoperative features, produces the highest area under the ROC value (0.920, 95%CI 0.907-0.932), although the difference is not statistically significant as compared with the imputed value. Reducing the preoperative feature set by removing variables indicating when the lab tests were administered before surgery further increases the area under the ROC of both the preoperative features and imputed-ASA (0.921, 95%CI 0.908-0.934) and the preoperative features and true ASA status (0.929, 95%CI 0.915-0.943). The Random Forest model has the best performance of all models used (see Table 2).

**Table 2.** Mortality Prediction Performance using AUROC curve.

| Model / AUC (95% CI) | Logistic Regression | ElasticNet Classifier | Random Forest | XGBoost Classifier |
| --- | --- | --- | --- | --- |
| **Charlson Comorbidity** | **0.706**<br>(0.669-0.744) | **0.706**<br>(0.669-0.744) | 0.692<br>(0.655-0.728) | 0.692<br>(0.655-0.728) |
| **ASA Status** | **0.865**<br>(0.848-0.882) | **0.865**<br>(0.848-0.882) | 0.828<br>(0.766-0.890) | 0.828<br>(0.766-0.890) |
| **Preop Features** | 0.872<br>(0.840-0.903) | 0.888<br>(0.860-0.917) | **0.909**<br>(0.881-0.937) | 0.895<br>(0.863-0.927) |
| **Preop + ASA Status** | 0.883<br>(0.855-0.910) | 0.905<br>(0.883-0.928) | **0.920**<br>(0.907-0.932) | 0.919<br>(0.904-0.934) |
| **Preop + imputed-ASA** | 0.868<br>(0.841-0.895) | 0.890<br>(0.868-0.912) | **0.912**<br>(0.902-0.923) | 0.901<br>(0.884-0.917) |
| **Preop (No Time) + ASA Status** | 0.886<br>(0.859-0.914) | 0.908<br>(0.883-0.932) | **0.929**<br>(0.915-0.943) | 0.916<br>(0.904-0.927) |
| **Preop (No Time) + imputed-ASA** | 0.871<br>(0.841-0.900) | 0.891<br>(0.868-0.914) | **0.921**<br>(0.908-0.934) | 0.897<br>(0.881-0.912) |

Area under the ROC (AUROC) curve values for each model and each of the seven input feature sets, using 10-fold cross-validation. Models with the highest AUROC are shown in bold. The mean value of the AUROC is shown, along with the 95% confidence interval ( $\bar{x} \pm 1.96 \times \sigma/\sqrt{n}$ ) in parenthesis. When using either the Charlson comorbidity or the ASA status as the only input feature, the linear models (Logistic regression, ElasticNet) outperform the non-linear models (Random Forest, XGBoost). However, for feature sets using more than one feature, the non-linear models outperform the linear models. In particular, the Random Forest has the highest AUROC compared to the other models.

**Fig 2.**
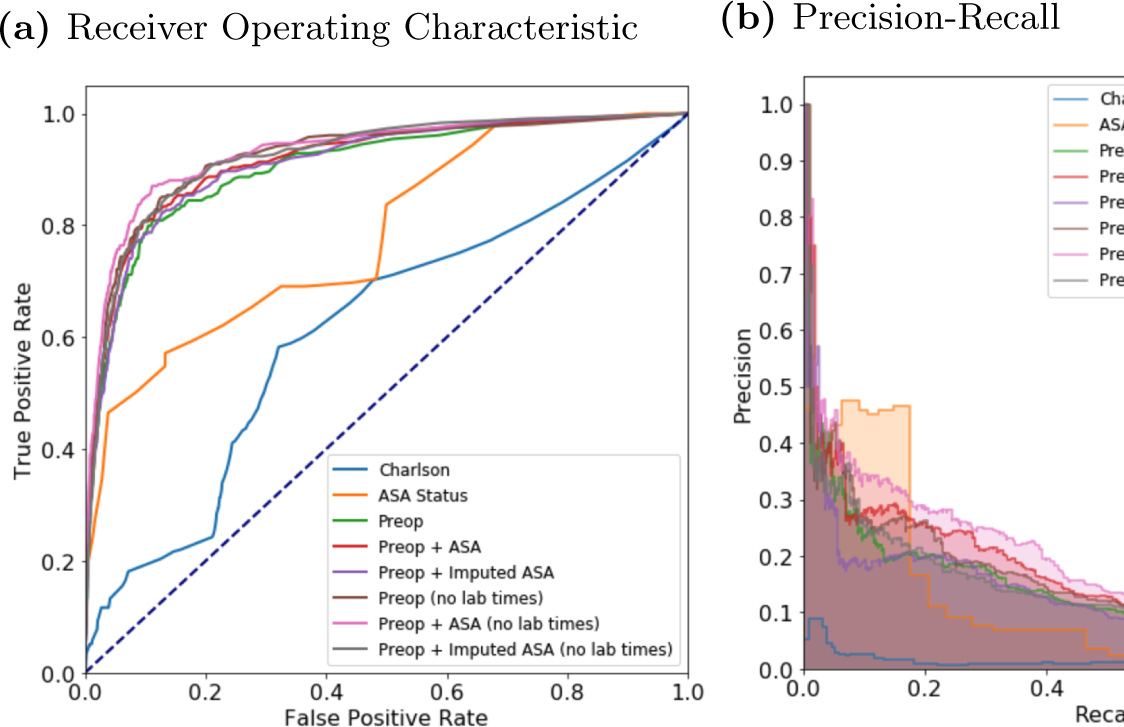
Receiver operating characteristic (ROC) and precision recall curves for the random forest model. Plots were generated using 10-fold cross-validated predictions on the entire dataset. ROC curves (a) show the false positive rate on the x-axis and the true positive rate on the y-axis. The optimal point is the upper-left corner. Precision-recall curves (b) show the recall on the x-axis and precision on the y-axis. The optimal point is in the upper-right corner.

#### 2.2.2 Calibration

A well-calibrated binary classification model outputs probabilities that are close to the true label (either a 1 for positive examples, or 0 for negative examples). Model calibration is often measured using the Brier score, which is the average squared distance between the predicted probability of the outcome and the true label over all samples. We used this metric to assess the calibration of our models, as shown in Table 3. The non-linear models (Random Forest, XGBoost) had much lower Brier scores compared to the linear models (Logistic Regression, ElasticNet). When using either the Charlson comorbidity score or the ASA status as the only feature, the XGBoost classifier had the lowest Brier score (0.0468 and 0.0486, respectively). For the the other five feature sets the random forest models obtained the lowest Brier scores (0.0060, 0.0056, 0.0058, 0.0055, and 0.0057, respectively).

**Table 3.**
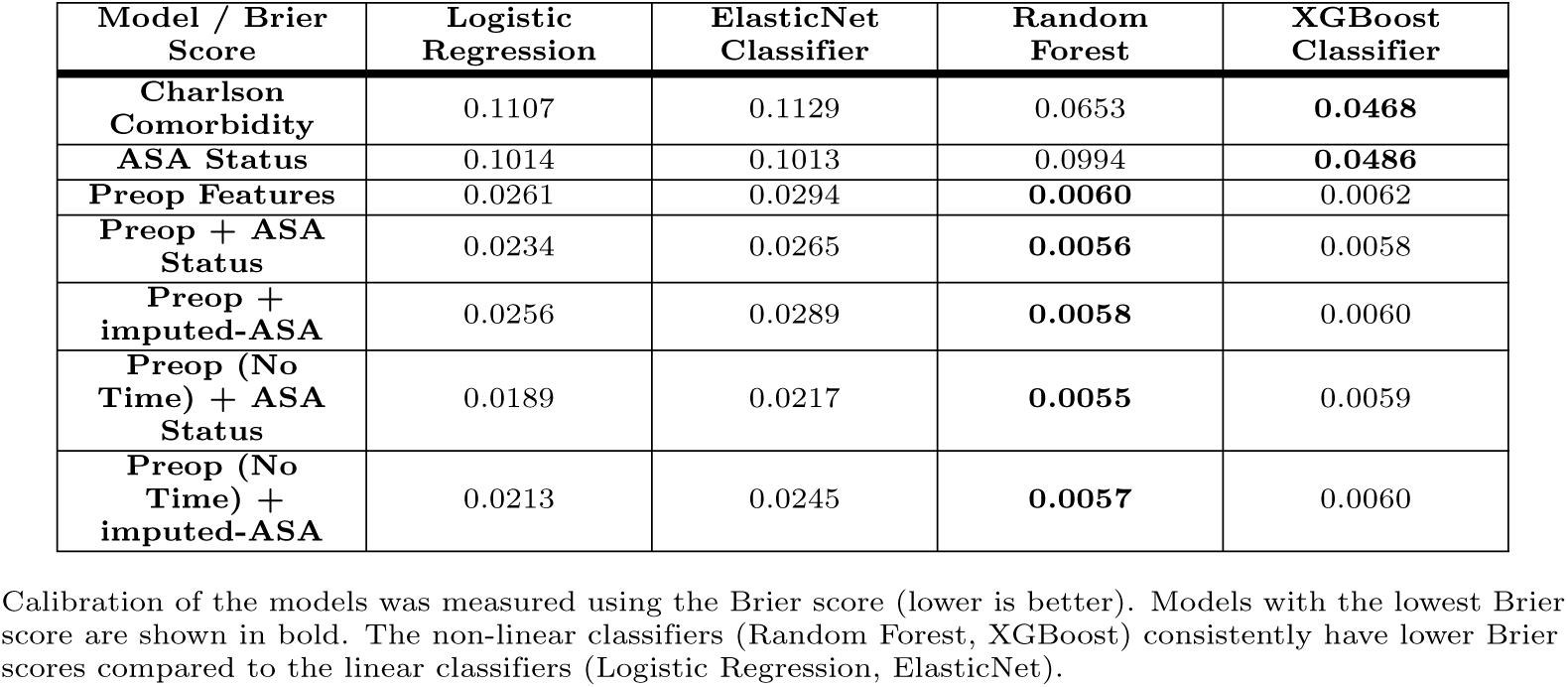
Calibration of Models: Brier Score.

#### 2.2.3 Precision-Recall

While the ROC curves are very informative of binary classification prediction performance in general, the precision-recall curves can be more informative when the classes are highly imbalanced [30]. However, these plots are often not included in methods for predicting surgical mortality, in which the datasets are inherently imbalanced.

An optimal model would reach the point in the upper-right corner of the PR plot (i.e. perfect recall and perfect precision). Using the random forest model, PR curves for each of the sets of features are shown in Figs 2b and, for all models, in Fig S2. As shown in the plots, models trained with a highly imbalanced dataset often suffer from poor precision-recall. Including the ASA status as an input feature increases the precision-recall compared to only using the preoperative features alone. The feature set using only the ASA status does surprisingly well, likely due to individuals with an ASA score of 5 being highly enriched for mortality and the difficulty of imputing ASA scores.

A clinically-relevant question to ask is how many additional patients must be flagged to capture one additional mortality; this is an extension of the “number needed to treat” idea for a risk ordered population. In Fig 3, the marginal increase in the number needed to be flagged is plotted as a function of the number of mortalities captured, using the risk scores generated from the random forest model. For the first hundred mortalities, this number ranges between one and ten with limited exceptions. After the first hundred mortalities however, the marginal increase is often in the hundreds and sometimes thousands of additional patients.

**Fig 3.**
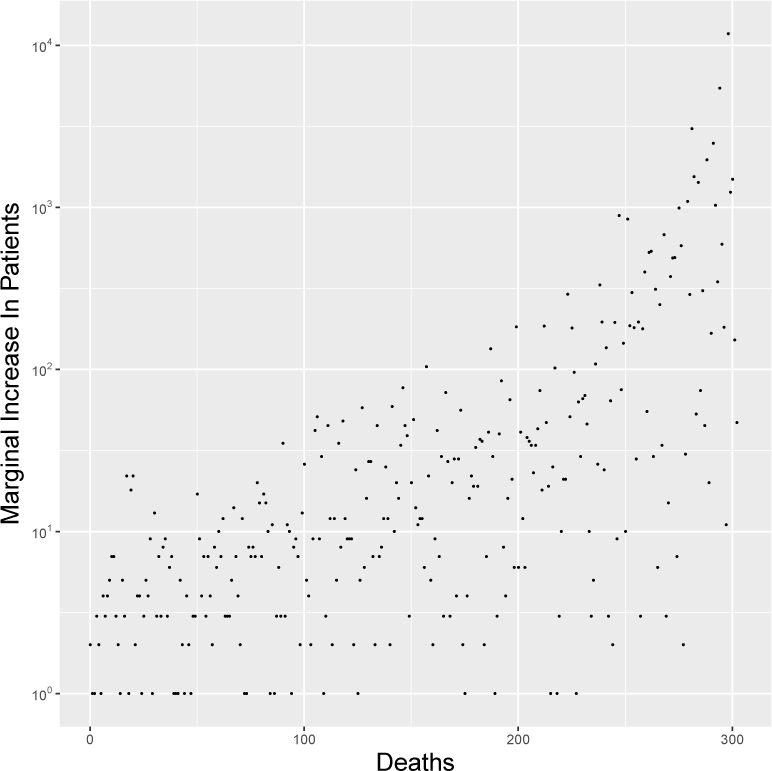
Marginal increase in number of patients needed to capture one additional death. This figure shows the marginal increase in the number of patients that need to be treated as a function of the number of in-hospital mortalities that will be captured. Note that the marginal increase in number of patients is log-scaled.

### 2.3 Feature Importance

To determine the most important features for each of the models, we examined the feature weights of the linear models and feature importances of the non-linear models. The top five most important features of each model, when trained on the preoperative features excluding the time to surgery of laboratory tests, are shown in Table 6. For the feature importance using other combinations of ASA and time to surgery of laboratory tests, see Tables S10, S11 and S12. As expected [20–22, 31], the ASA status is the most important feature of every model where it is contained. Interestingly, there were as many important features shared by the ElasticNet model and the XGBoost model as there were by the random forest and the XGBoost model (as shown by the bold numbers in Table 6). Notably, the random forest model placed more importance on surgery-specific information, like patient class and pre-surgery location, whereas the XGBoost model placed higher importance on the lab results.

In Table 7, the feature importance for the random forest model is shown using four different sets of input features. For the feature sets that include lab result timestamps, many of the most important features are the lab result time-stamp features. However, when these features are removed, the feature importance shifts to the lab results themselves, as well as surgery-specific features such as the patient class.

### 2.4 Integrating Preoperative Risk with Postoperative Risk

Replacing the ASA status with the preoperative risk predictions in the postoperative risk prediction model generated similar results. The postoperative risk model that was trained using the preoperative risk scores had an area under the ROC of 0.924 (95%CI 0.905-0.941), whereas the postoperative model trained using the ASA status had an area under the ROC of 0.929 (95%CI 0.911-0.944). This is in line with the previously published results of this model [11]. Interestingly, the area under the ROC of the preoperative model is similar to the area under the ROC of the postoperative model, which suggests that the postoperative features may not be capturing additional information.

In order to examine how mortality risk changes from immediately before surgery to after surgery, the preoperative and postoperative risk scores for all patients were grouped by percentiles and the counts of each grouping are displayed in Table 4 and with a heatmap (Fig 4a). For the majority of patients we see a slight increase or decrease in their postoperative risk compared to the initial preoperative risk, as demonstrated by the coloring just above/below the diagonal line in Fig 4a. Fig 4b demonstrates the same plot, but contains only those patients who eventually died during that admission, and the corresponding data can be found in Table 5. Note that most of these patients fall in the upper right quadrant high-risk quadrant of the heatmap (Fig 4b). Moreover, 78% of patients who die and have a preoperative risk percentile below 95% have an increased postoperative risk percentile. This is substantially greater than the percent of matched patients from a null distribution who have an increased percentile (see Fig S5).

**Table 4.**
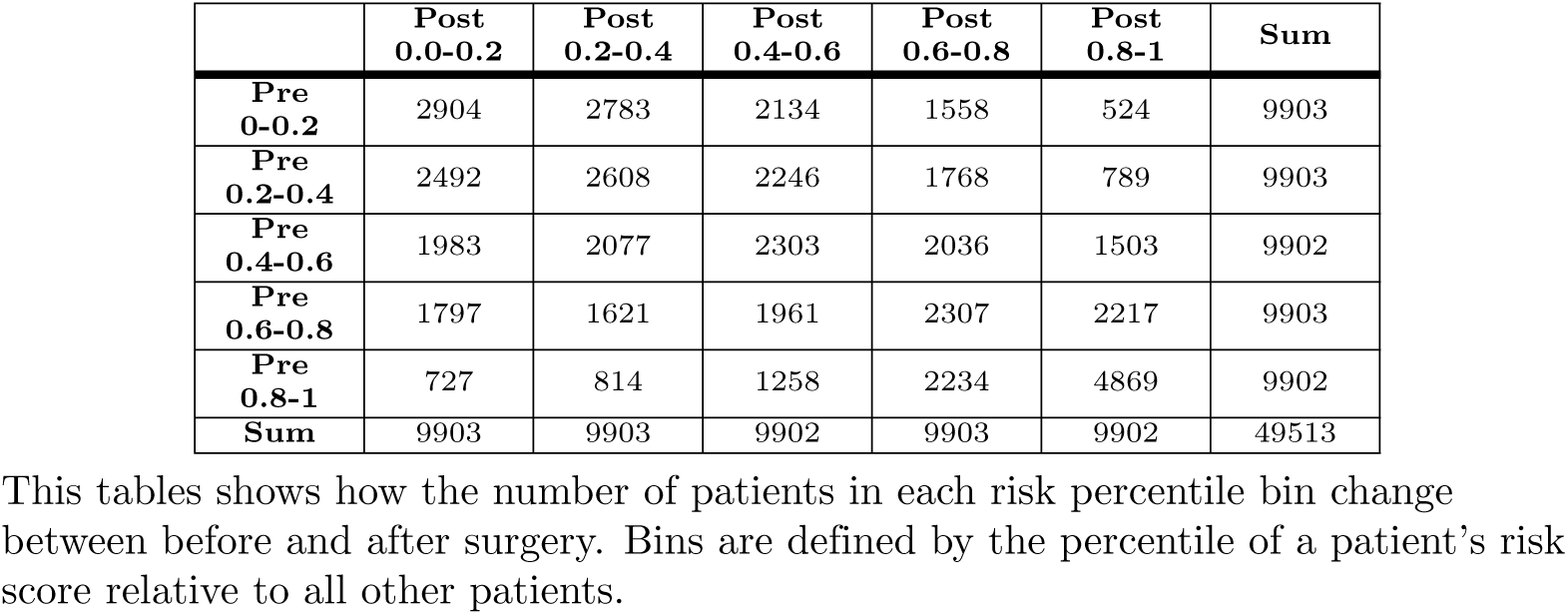
Flow of from Pre- to Post-operative Percentile Bins.

**Table 5.** Change in Pre- to Post-operative Risk Percentile Bins for In-hospital Mortalities.

|  | Post<br>0.0-0.2 | Post<br>0.2-0.4 | Post<br>0.4-0.6 | Post<br>0.6-0.8 | Post<br>0.8-1.0 | Sum |
| --- | --- | --- | --- | --- | --- | --- |
| **Pre**<br>**0-0.2** | 0 | 0 | 0 | 0 | 1 | 1 |
| **Pre**<br>**0.2-0.4** | 0 | 0 | 0 | 3 | 5 | 8 |
| **Pre**<br>**0.4-0.6** | 0 | 0 | 1 | 4 | 6 | 11 |
| **Pre**<br>**0.6-0.8** | 0 | 0 | 1 | 0 | 20 | 21 |
| **Pre**<br>**0.8-1** | 4 | 0 | 5 | 9 | 244 | 262 |
| **Sum** | 4 | 0 | 7 | 16 | 276 | 303 |

This tables shows how the number of in-hospital mortalities in each risk percentile bin change between before and after surgery. There is an increase in the number of postoperative high-risk patients (0.8-1.0) compared to preoperative high-risk patients (0.8-1.0).

**Table 6.** Model Feature Importance: Without ASA and Without Lab Time Features.

| Feature | Random Forest | Linear Regression | Elastic Net | XGBoost |
| --- | --- | --- | --- | --- |
| ALBUMIN | **0.04\*** | -0.57 | -0.48 | **0.03\*** |
| BNP | **0.04\*** | **1.45\*** | **1.12\*** | **0.03\*** |
| HEMOGLOBIN | **0.05\*** | -0.8 | -0.77 | **0.03\*** |
| INR | **0.05\*** | 1.14 | 0.76 | 0.02 |
| PAT CLASS INPATIENT | **0.06\*** | -0.67 | -0.12 | 0.02 |
| PAT CLASS SAME DAY ADMIT | **0.04\*** | -1.31 | -0.51 | 0 |
| PLATELET.COUNT | **0.04\*** | -0.89 | **-0.92\*** | **0.04\*** |
| PRE SURG LOCATION PRE-ADMISSION | **0.05\*** | -0.38 | -0.66 | 0.01 |
| DIAS.BLOOD.PRESSURE | 0.03 | -0.9 | **-0.82\*** | **0.03\*** |
| HCUP CODE 39.0 | 0 | 1.08 | **0.79\*** | 0 |
| SPO2 | 0.02 | **-3.85\*** | **-0.97\*** | **0.03\*** |
| AGE LT 89 | 0.01 | 1.02 | 0.55 | **0.03\*** |
| ALKALINE.PHOSPHATASE | 0.01 | 0.26 | 0.02 | **0.03\*** |
| ALT | 0.01 | -0.42 | -0.11 | **0.03\*** |
| AST | 0.01 | 0.85 | 0.4 | **0.03\*** |
| BICARBONATE | 0.03 | 0.31 | 0.04 | **0.03\*** |
| BILIRUBIN.TOTAL | 0.02 | 0.16 | 0.35 | **0.03\*** |
| BMI | 0.01 | 0.17 | 0.12 | **0.04\*** |
| ECHO.EF | 0.02 | -0.82 | -0.5 | **0.03\*** |
| GLUCOSE | 0.02 | 0.89 | 0.63 | **0.03\*** |
| HEIGHT IN | 0.01 | **-1.64\*** | -0.62 | **0.03\*** |
| HRS ADMSN TO SURGERY | 0.03 | 0.69 | 0.57 | **0.04\*** |
| POTASSIUM | 0.01 | -0.73 | -0.66 | **0.03\*** |
| PROTHROMBIN.TIME | 0.03 | 0.46 | 0.2 | **0.03\*** |
| PULSE | 0.01 | 0.31 | 0.17 | **0.03\*** |
| SYS.BLOOD.PRESSURE | 0.02 | -0.35 | -0.2 | **0.04\*** |
| UREA.NITROGEN | 0.02 | 0.09 | 0.12 | **0.03\*** |
| WHITE.BLOOD.CELL.COUNT | 0.01 | 0.76 | 0.65 | **0.03\*** |
| CASE SRV NAME Otolaryngology | 0 | **-1.5\*** | -0.43 | 0 |
| CHLORIDE | 0.01 | **1.85\*** | 0.3 | 0.02 |

**Table 7.** Random Forest Feature Importance.

| Feature | With ASA<br>and Lab<br>Times | With ASA<br>and No Lab<br>Times | No ASA and<br>With Lab<br>Times | No ASA and<br>No Lab Times |
| --- | --- | --- | --- | --- |
| ASA STATUS | **0.07\*** | **0.08\*** | NA | NA |
| PRE SURG<br>LOCATION<br>PRE-ADMISSION | **0.04\*** | **0.05\*** | **0.04\*** | **0.05\*** |
| PAT CLASS<br>INPATIENT | **0.04\*** | **0.05\*** | **0.04\*** | **0.06\*** |
| UREA.NITROGEN.HRS<br>2 SURGERY | **0.03\*** | NA | **0.03\*** | NA |
| POTASSIUM.HRS 2<br>SURGERY | **0.03\*** | NA | 0.02 | NA |
| PLATELET.COUNT.HRS<br>2 SURGERY | **0.03\*** | NA | **0.03\*** | NA |
| HEMOGLOBIN.HRS 2<br>SURGERY | **0.03\*** | NA | 0.02 | NA |
| HEMOGLOBIN.HRS 2<br>SURGERY | **0.03\*** | NA | 0.02 | NA |
| CREATININE.HRS 2<br>SURGERY | **0.03\*** | NA | **0.03\*** | NA |
| CHLORIDE.HRS 2<br>SURGERY | **0.03\*** | NA | 0.02 | NA |
| HEMOGLOBIN | 0.01 | 0.03 | 0.01 | **0.05\*** |
| INR | 0.02 | **0.04\*** | **0.03\*** | **0.05\*** |
| ALBUMIN | 0.01 | **0.04\*** | 0.02 | **0.04\*** |
| BNP | 0.01 | 0.03 | 0.01 | **0.04\*** |
| PLATELET.COUNT | 0.01 | 0.03 | 0.01 | **0.04\*** |
| PAT CLASS SAME<br>DAY ADMIT | 0.02 | **0.04\*** | **0.03\*** | **0.04\*** |

The top five features of the random forest model, ranked by Gini importance, for each feature set are shown in bold and have an asterisk. Features with a NA value were not included in the dataset. All models included the preoperative features. “No ASA” means the ASA status was not included as a feature, and “No Lab Times” means the timestamps of the lab results were not included (i.e. features with the “.HRS\_2.SURGERY” suffix).

**Fig 4.**
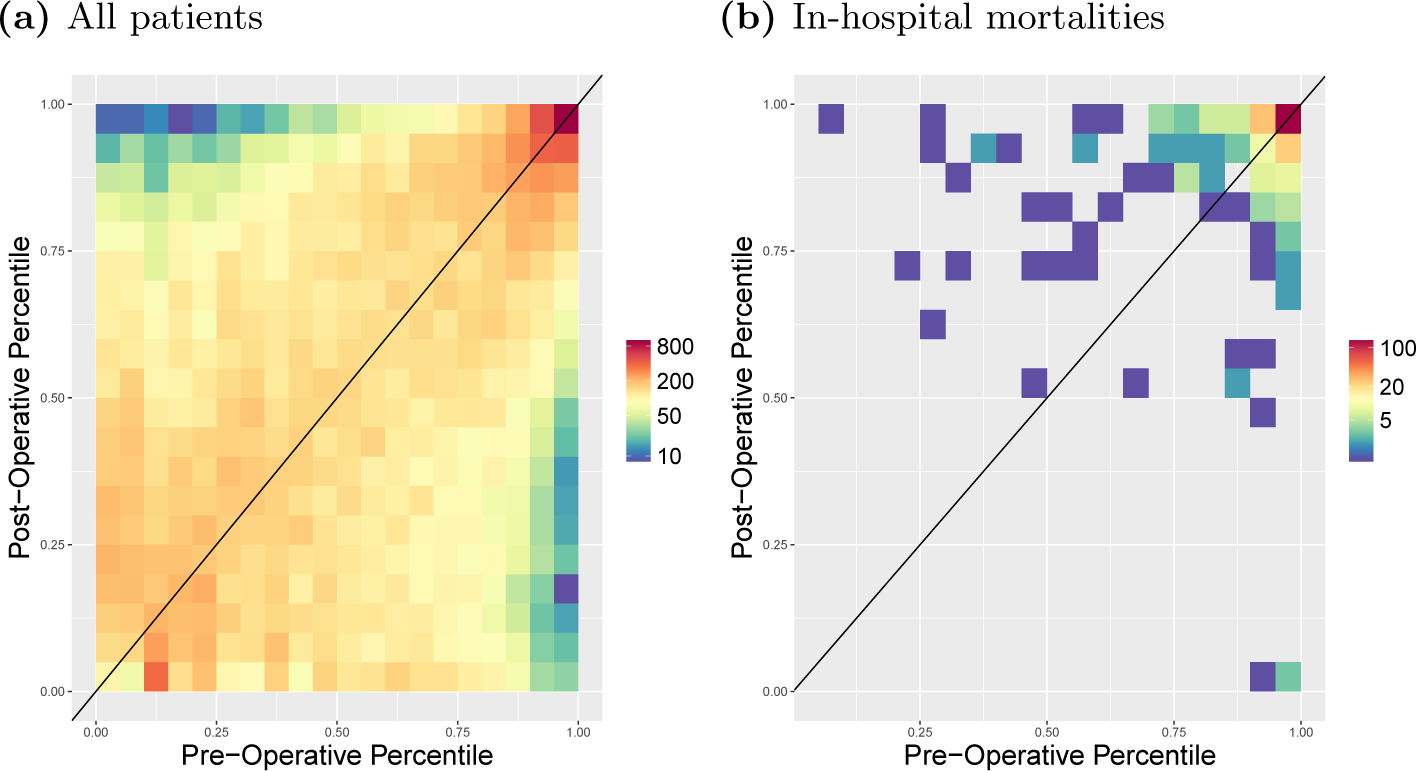
Heatmap of Preoperative Risk vs. Postoperative Risk. Preoperative (x-axis) and postoperative (y-axis) risk scores were binned by percentile, and the counts per bin visualized as a heatmap in log scale. Preoperative risk predictions were generated using the random forest model trained on the preoperative features, including lab times, and the imputed-ASA status. In (a) all patients are displayed, and in (b) only the in-hospital mortalities are shown. 78% of patients who die and have a pre-operative risk percentile below 95% have an increased postoperative risk percentile. This is substantially greater than the percent of matched patients from a null distribution who have an increased percentile, see Fig S5.

The change in risk between the preoperative and postoperative time points is shown using violin plots in Fig 5, where the risk scores are binned by percentile and stratified by ASA status. In all four subplots of Fig 5, the in-hospital mortalities are, on average, at higher risk than the patients who survive surgery, as shown by the increased mass toward the top of the blue plots compared to the red plots. This is particularly apparent in Figs 5a and 5b as the majority of patients have an ASA status of 2 and 3, and therefore the violins are larger. In Figs 5c and 5d, most of the patients begin and end surgery at high risk in both the preoperative model and postoperative model, and we can see that the models correctly rank those that die at higher risk, as shown by the increased mass toward the top of the blue plots compared to the red. Violin plots for patients with an ASA status of 1 are not shown as none of those patients died during the admission.

**Fig 5.**
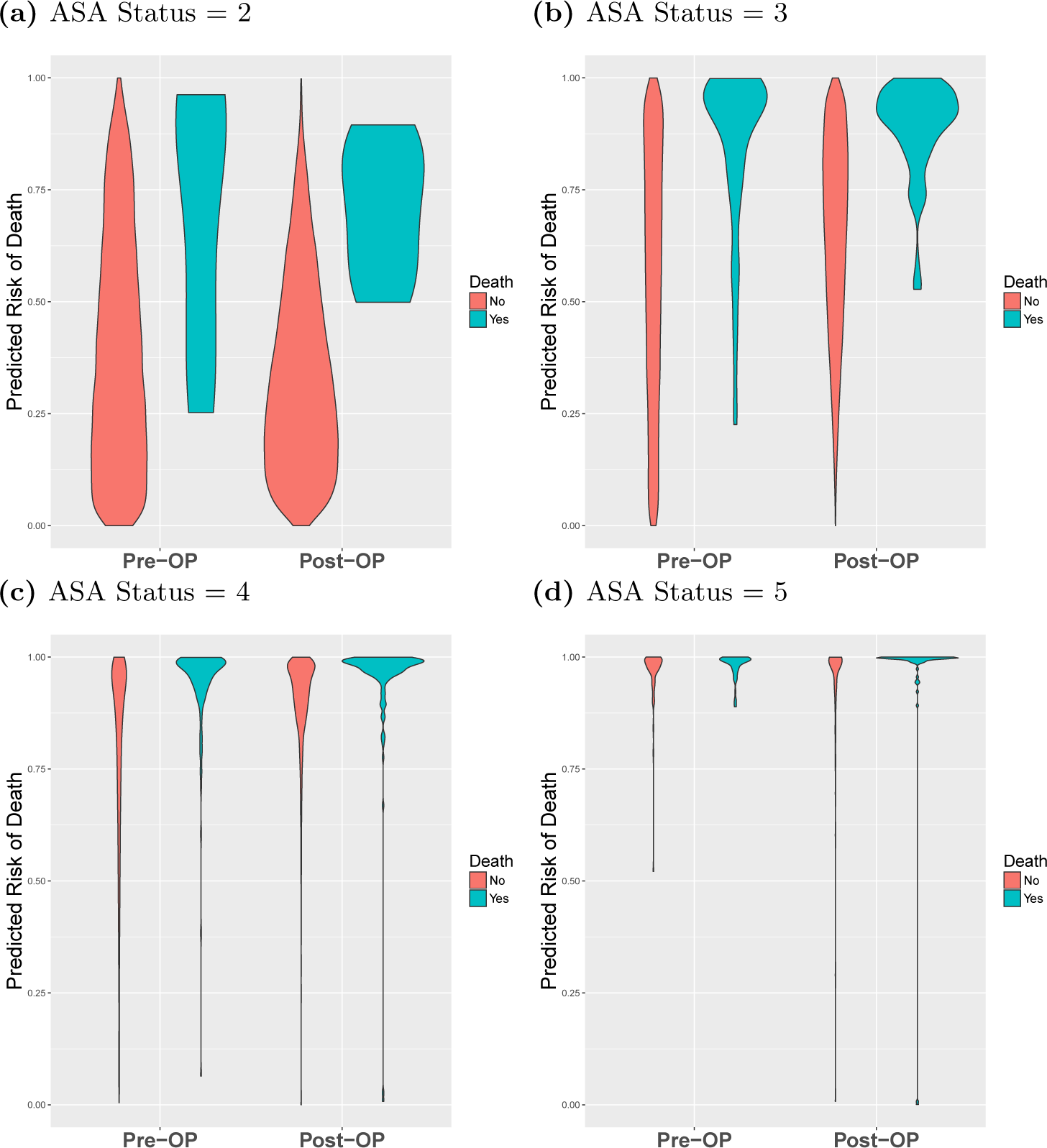
Comparison of preoperative and postoperative mortality predictions binned by percentile. Violin plots showing the percentile rankings of the mortality risk prediction scores relative to all other individuals, stratified by ASA status. Preoperative risk (2 left violins of each plot) and postoperative risk (right 2 violins of each plot) are shown in each plot. Red violins are patients that survived surgery, and blue violins are mortalities.

## 3 Discussion

In this manuscript we were able to successfully create a fully-automated preoperative risk prediction score that can better predict in-hospital mortality than both the ASA Score and the Charlson comorbidity index. Unlike the ASA status or the Charlson comorbidity scores, this score was built using solely objective clinical information that was readily available from the EMR prior to surgery. Similar to previous models [11], the results indicate that inclusion of the ASA score in the model improves the predictive ability. However, we were able to account for this by “imputing” the ASA score and including this additional feature in our model. We were additionally able to integrate the results of our model into a previously developed postoperative risk prediction model and achieve a performance that was comparable to the use of the ASA physical status score in that model. Lastly, when using the preoperative and postoperative scores together we were able to demonstrate that, on a patient level, risk does change during the perioperative period - indicating that choices made in the operating room may have profound implications for our patients.

The challenge of perioperative risk stratification is certainly not new. In fact, the presence of so many varied risk scores (ASA Score, Charlson Comorbidity Index [6], POSPOM [4], RQI [32], NSQIP Risk Calculator [8]) speaks to the importance with which clinicians view this problem. A major limitation of many of these models has been that they either rely on data not available at the time of surgery (i.e. ICD codes), or they require an anesthesiologist to review the chart (those that contain the ASA score). The fact that our model can be fully automated and yet perform better than these models implies that it has broad applicability.

As demonstrated in many previous studies [17, 20–22], the ASA score itself remains a good predictor of postoperative outcomes. We believe that one reason that our model is able to outperform the ASA score is related to the overwhelming amount of information contained in the modern patient’s medical record. The introduction of the EMR has lead to an explosion of information that can frequently make it challenging for a clinician to consume everything. For example, due to time constraints, an anesthesiologist may not be able to fully assess the impact of every entry in a patient’s health record before an upcoming procedure. This limitation of a clinician’s ability to find all of the relevant information in the modern EMR highlights the advantage of an automated scoring system such as this - not as a replacement for physicians - but rather as a tool to help them better focus their efforts on those patients most likely to benefit.

Our efforts to leverage the value of the ASA score led us to create an “imputed” ASA score for us to include in our model. However, the creation and inclusion of this feature did not significantly improve the predictive power of our model. It is important to note that this is not truly an imputation of, or replacement to, the ASA score. Our techniques and the ASA score differ in some key respects. First, while the ASA score is an ordinal score with values between 1 and 5, our score is actually a continuous variable with no such thresholds. Second, the ASA score itself is not designed to predict postoperative mortality *per se*, but rather is a set of clinical guidelines that are used by practicing clinicians to categorize their patients. We believe that the fact that the ASA score is not designed to predict any specific postoperative outcome is a potential advantage of using these analytic techniques. While the model that is presented here was optimized to focus on postoperative mortality, the same techniques could easily be used to help predict a wide range of postoperative outcomes such as acute kidney injury, respiratory failure, readmission and others. Such a refined classification would not be possible with the ASA or any other generalizable risk score.

Another advantage of an automated model such as this is that it allows for the continuous recalculation of risk longitudinally over time. As shown in Fig 4, most patients have either a minor increase or decrease in risk in the time from before to after surgery and, unsurprisingly, in patients who eventually die, that risk tends to increase. We believe that this finding has several important implications. First, given the challenges of continually monitoring the risk of all patients in the hospital, advanced analytical models such as the models proposed in this manuscript have great potential to act as early warning systems alerting clinicians to sudden changes in risk profiles and facilitating the use of rapid response teams. Second, the frequency with which risk changed substantially during the operative period highlights the effect to which intraoperative interventions may have implications far beyond the operating room. Multiple specific interventions, including the avoidance of intraoperative hypotension and hypothermia, have been shown to have effects on longer-term outcomes, and currently enhanced recovery after surgery (ERAS) pathways have promoted the standardization of intraoperative interventions. We believe that our findings should add to the evidence that a well-prescribed anesthetic plan may be of longer-term benefit to patient outcomes.

One potential promise of the use of machine learning in medicine is the ability to leverage these models in order to better understand what features are truly driving outcomes. In an effort to better understand this, we extracted the weights of the features in both the linear and non-linear models. Of note, removing some features, specifically the relative time of lab tests, actually improved the results of our model. This could potentially be caused by multiple correlated features tagging an underlying cause, and the correlation introduces noise in the model as the importance is distributed among multiple features rather than focused on a single feature. In theory, a machine learning model should be able to remove these features by setting a coefficient to zero. However, in practice, this may not always be the case - as illustrated here, where we force this behavior by manually removing the features from the model. We believe that this finding highlights the importance of having collaborative relationships between experts in machine learning and clinicians who are able to help guide which features to include in a model. Simply entering large amounts of data from an EMR, without proper clinical context, is unlikely to create the most effective or efficient models.

There are several key limitations of this study. Perhaps the most significant is the low frequency of the outcome in question - in-hospital mortality. The incidence of mortality was less than 1% - implying that a model that blindly reports “survives” every time will have an accuracy greater than 99%. Predicting such a rare outcome makes it highly challenging to produce results with very high precision. Nonetheless, the models presented in this paper do outperform other models currently in use, as measured by area under the ROC curve. Secondly, the data used here are from a single large academic medical center. Thus, it is possible, though unlikely, that this model will not perform similarly at another institution. Past experience has demonstrated that, with recalibration, models can easily be transported from one institution to another. One last limitation lies not necessarily with the study itself but rather with the overall landscape of EMR data. While the promises of fully-automated risk scores are great, the reality remains that most institutions still have trouble accessing the data stored in the EMRs. Thus, in order to truly automate processes such as these, robust data interoperability standards (such as Fast Healthcare Interoperability Resources (FHIR)) will be needed in order to allow access to data.

## 4 Conclusion

The promise of using machine learning techniques in healthcare is great. In this work we have presented a novel set of easily-accessible (via EMR data) preoperative features that are combined in a machine learning model for predicting in-hospital post-surgical mortality, which outperforms current clinical risk scores. We have also shown that the risk of in-hospital mortality changes over time, and that monitoring that risk at multiple time-points will allow clinicians to make better data-driven decisions and provide better patient care. It is our hope and expectation that the next few years will produce a plethora of research leveraging data obtained during routine patient care to improve care delivery models and outcomes for all of our patients.

## Supporting Information

**Fig S1. Receiver Operating Characteristic (ROC) curves** ROC curves show the false positive rate on the x-axis and the true positive rate on the y-axis. The optimal point is the upper-left corner. Plots were generated using cross-validated predictions on entire dataset. Models that use preoperative features consistently outperform models that only use Charlson score or ASA status. Logistic Regression (a) and ElasticNet (b) (linear models) outperform Random Forest (c) and XGBoost (d) (non-linear models) when using a single input feature. However, non-linear models (c, d) outperform linear models (a, b) when using multiple features.

**Fig S2. Precision-recall (PR) curves** PR curves show the recall on the x-axis and precision on the y-axis. The optimal point is in the upper-right corner. Figs (a) and (b) show that the linear models have very similar PR curves. The gradient boosted trees model (d) has better precision-recall compared to the random forest (c).

**Fig S3. Change in preoperative and postoperative mortality prediction rank** Points on the left of the plot are preoperative predictions of mortality, and points on the right are postoperative predictions of mortality. The preoperative probability predictions were ranked from highest to lowest, and for each patient a line from the preoperative prediction is then connected to the rank in list of sorted postop predictions. If a point does not change its rank in the list of risk scores, then the line will be straight across. Otherwise if the rank increased, it will have positive slope, or negative slope if it decreased. Blue lines are patients that survived surgery, and red lines are in-hospital mortalities. The median survivor points are displayed by the yellow line. Spearman rank-order correlation coefficient and p-value are shown above each plot.

**Fig S4. Comparison of preoperative and postoperative mortality prediction using risk scores** Violin plots showing the distributions of mortality risk, stratified by ASA status. Preoperative risk (2 left violins of each plot) and postoperative risk (right 2 violins of each plot) are shown in each plot. Red violins are patients that died during surgery, and blue violins are survivors.

**Fig S5. Null distribution of increasing risk for matched individuals based on similar pre-operative scores** For each individual who died, we found a matched individual who falls within one percentile of the pre-operative risk percentile of the individual that died. We then calculated the number of individuals in this matched set who had increased chance of death for their post-operative score. We repeated this process 10,000 times. This histogram represents the percent of matched individuals that have a higher post-operative score than pre-operative score. We restrict our analysis to individuals who fall in less than the ninety-fifth percentile.

**Table S1. Feature List** Preoperative features used in the model. All features are readily available via the EHR system prior to surgery. HCUP codes are included as features, but not shown in this table. Refer to Table S2 for a full list of HCUP codes.

**Table S2. Comparison of HCUP codes between overall surgeries and surgeries with death** The numbers in parentheses () in the Count and Percent columns represent the count and percentage of individuals who died in-hospital undergoing surgery who were classified under the given HCUP description.

**Table S3. Predicting Mortality using Preoperative Features** Model performance metrics for predicting mortality using preoperative features and 10-fold cross-validation.

True positives: TP, False positives: FP, True negatives: TN, False negatives: FN. Accuracy = (TP+TN)/(TP+TN+FP+FN). Precision = TP/(TP+FP).

Recall = TP/(TP+FN).

Specificity = TN/(TN+FP). F1 Score = 2/((1/Recall)+(1/Precision)).

**Table S4. Predicting Mortality using Preoperative Features + ASA Status** Model performance metrics for predicting mortality using both preoperative features and ASA status as input features, and 10-fold cross-validation. For an explanation of the metrics see the description of Table S3

**Table S5. Predicting Mortality using ASA Status** Model performance metrics for predicting mortality using only the ASA status as an input feature, and 10-fold cross-validation. For an explanation of the metrics see the description of Table S3

**Table S6. Predicting Mortality using Charlson Comorbidity** Model performance metrics for predicting mortality using only the Charlson comorbidity as an input feature, and 10-fold cross-validation. For an explanation of the metrics see the description of Table S3

**Table S7. Predicting Mortality using Preoperative Features + imputed-ASA Score** Model performance metrics for predicting mortality using both preoperative features and the imputed-ASA score as input features, and 10-fold cross-validation. For an explanation of the metrics see the description of Table S3

**Table S8. Predicting Mortality using Preoperative Features + ASA status, Without Lab Times** Model performance metrics for predicting mortality using both preoperative features and the ASA score but without lab times as input features, and 10-fold cross-validation. For an explanation of the metrics see the description of Table S3

**Table S9. Predicting Mortality using Preoperative Features + imputed-ASA status, Without Lab Times** Model performance metrics for predicting mortality using both preoperative features and the imputed-ASA score but without lab times as input features, and 10-fold cross-validation. For an explanation of the metrics see the description of Table S3

**Table S10. Model Feature Importance: With ASA and Without Lab Time Features** The bold values with the * indicates that the value was one of the top five strongest predictors for the given model. In the Logistic Regression and ElasticNet columns the model weights are shown, where negative values indicate the feature lowers the probability of mortality. The non-linear model features are ranked according to the mean decrease in Gini impurity.

**Table S11. Model Feature Importance: Without ASA and With Lab Time Features** The bold values with the * indicates that the value was one of the top five strongest predictors for the given model. In the Logistic Regression and ElasticNet columns the model weights are shown, where negative values indicate the feature lowers the probability of mortality. The non-linear model features are ranked according to the mean decrease in Gini impurity.

**Table S12. Model Feature Importance: With ASA and Time Features** The bold values with the * indicates that the value was one of the top five strongest predictors for the given model. In the Logistic Regression and ElasticNet columns the model weights are shown, where negative values indicate the feature lowers the probability of mortality. The non-linear model features are ranked according to the mean decrease in Gini impurity.

**Table S13. Change in Pre- to Post-operative Risk using ASA-like risk bins** This tables shows how risk changes for groups defined by the number of patients with each ASA score, from 1 to 5. Patient risk scores were sorted from highest-risk to lowest risk, and then patients were binned into 5 categories, where the number of patients in each category corresponds to the number of patients in each ASA status group. For example, 127 patients had an ASA status of 5, therefore, bins Pre 5 and Post 5 each have 127 patients.

**Table S14. Change in Pre- to Post-operative Risk using ASA-like risk bins for In-hospital Mortalities** This tables shows how risk changes for groups defined by the number of patients with each ASA score, from 1 to 5. Patient risk scores were sorted from highest-risk to lowest risk, and then patients were binned into 5 categories, where the number of patients in each category corresponds to the number of patients in each ASA status group. In this table we only show in-hospital mortalities. The number of patients in group 5 increases between the preoperative and postoperative periods, showing that the postoperative model is capturing the increased risk derived from intraoperative data.

## Acknowledgments

L.O.L. was financially supported by the National Institute of Mental Health under award number K99MH116115. S.S. was supported in part by NIH grants R00GM111744, R35GM125055, NSF Grant III-1705121, an Alfred P. Sloan Research Fellowship, and a gift from the Okawa Foundation. R.B. was funded by a UCLA QCB Collaboratory Postdoctoral Fellowship directed by Matteo Pellegrini. U.M. was financially supported by the Bial Foundation. B.J. was supported by the National Science Foundation Graduate Research Fellowship Program under Grant No. DGE-1650604.

M.C. is co-owner of US patent serial no. 61/432,081 for a closed-loop fluid administration system based on the dynamic predictors of fluid responsiveness which has been licensed to Edwards Lifesciences. M.C. is a consultant for Edwards Lifesciences (Irvine, CA), Medtronic (Boulder, CO), Masimo Corp. (Irvine, CA). M.C. has received research support from Edwards Lifesciences through his Department and NIH R01 GM117622 - Machine learning of physiological variables to predict diagnose and treat cardiorespiratory instability and NIH R01 NR013912 - Predicting Patient Instability Non invasively for Nursing Care-Two (PPINNC-2). No funding bodies had 401 any role in study design, data collection and analysis, decision to publish, or 402 preparation of the manuscript.

